# Electrophysiological correlates of syntactic structures

**DOI:** 10.1101/660415

**Authors:** Fiorenzo Artoni, Piergiorgio d’Orio, Eleonora Catricalà, Francesca Conca, Franco Bottoni, Veronica Pelliccia, Ivana Sartori, Giorgio Lo Russo, Stefano F. Cappa, Silvestro Micera, Andrea Moro

## Abstract

Syntax is traditionally defined as a specifically human way to pair sound with meaning: words are assembled in a recursive way generating a potentially infinite set of sentences^1,2^. There can be different phrasal structures depending on the types of words involved, for example, “noun phrases” (NP), combining an article and a noun, vs. “verb phrases” (VP), combining a verb and a complement. Although it is known that the combination of an increasing number of words in sequences correlates with an increasing electrophysiological activity^3,4^, the specific electrophysiological correlates of the syntactic operation generating NPs vs. VPs remain unknown. A major confounding factor is the fact that syntactic information is inevitably intertwined with the acoustic information contained in words even during inner speech^5^. Here, we addressed this issue in a novel way by designing a paradigm to factor out acoustic information and isolate the syntactic component. In particular, we construed phrases that have exactly the same acoustic content but that are interpreted as NPs or VPs depending on their syntactic context (homophonous phrases). By performing stereo-electro-encephalographic (SEEG) recordings in epileptic patients^6^ we show that VPs are associated with a higher activity in the high gamma band (150-300Hz frequency), an index of cortical activity associated with linguistic processing, with respect to NPs in multiple cortical areas in both hemispheres, including language areas and their homologous in the non-dominant hemisphere. Our findings pave the way to a deeper understanding of the electrophysiological mechanisms underlying syntax and contribute to the ultimate far reaching goal of a complete neural decoding of linguistic structures from the brain^2^.

---

Factoring sound out of a linguistic expression is a major empirical problem, since sound is inevitably intertwined with syntactic information^3,7^ even during inner speech^5^. By crucially relying on Italian grammar we designed a novel protocol to circumvent this problem and measure the specific electrophysiological correlates of two basic syntactic structures. The stimuli were pairs of different sentences containing strings of two words with exactly the same acoustic information but completely different syntax (homophonous strings). More specifically, each pair contained an NP, resulting from syntactic combination of two lexical elements (a definite article and a noun), and a VP, resulting from the syntactic combination of two different types of lexical elements (a verb and a pronominal complement): the NP and the VP were pronounced in exactly the same way. In addition, each VP included a further crucial difference: the object of the verb, realized as a pronoun, was moved from its canonical position on the right of the verb to the left of the verb, a syntactic operation called “cliticization”. For example, the sequence [la’pɔrta], could be interpreted either as a noun phrase (“the door”) or a verb phrase (“brings her”; lit.: her brings) depending on the syntactic context within the sentence where they were pronounced (Figure 1). As for the acoustic information concerning the homophonous phrases, it must be noticed that for each pair of sentences containing the same homophonous phrase, either phrase was deleted and substituted with a copy of the other one: this strategy was exploited to avoid the possibility that the structure of the two phrases could be distinguished by subtle intonational or prosodic clues: practically, the relevant part of the stimuli constituting the homophonous phrase was physically exactly the same. Although these results were based on a peculiar property of Italian language, our results are generalizable to other languages because the basic distinction of nouns vs. verbs is universally attested across-languages^8^. As for other variables constituting the homophonous phrases, words were balanced for major semantic features (such as abstract vs. concrete) and length (number of syllables). The two sentence types were further differentiated by the level of “surprisal”, an information-theoretic concept reflecting the expectedness of each word given its preceding context, which is defined as the negative log probability of a certain word in a sentence, given the words that precede it in that sentence^9^. The analysis shows that whereas there is no significant surprisal difference for the Verb/Noun position in the phrase, the values related to the article/clitic position were significantly different (Figure 2). In fact, the more complex syntactic structure, i.e. the VP involving movement of the object from the right to the left position of the verb, resulted in a higher surprisal level when the same auditory input was interpreted as a clitic rather than an article, as indicated by classical statistics and by decoding in the feature space with a Support Vector Machine (SVM) analysis. Extended Data Table 1 reports the number of valid cases, the percentage of missing, the mean and the standard deviation relative to the surprisal value, separately for the two experimental conditions. As reported in Extended Data Table 2, 84% (n = 26) of the sentences with low surprisal were NPs and 84% (n = 26) of the sentences with high surprisal were VPs. To fully exploit the potentials offered by homophonous strings we investigated the electrophysiological correlates of these NPs vs. VPs with intracranial electrodes for stereo-electro-encephalography (SEEG) monitoring (see Extended Data Figure 1 with the visualization information for one subject for the assessment of anatomical electrical sources). Invasive intracranial EEG monitoring allows us to precisely localize sources of activation. This procedure offers a unique opportunity to observe human brain activity with an unparalleled combination of spatial and temporal resolution, impossible otherwise by using the available non-invasive recording and imaging techniques.

**Figure 1.**
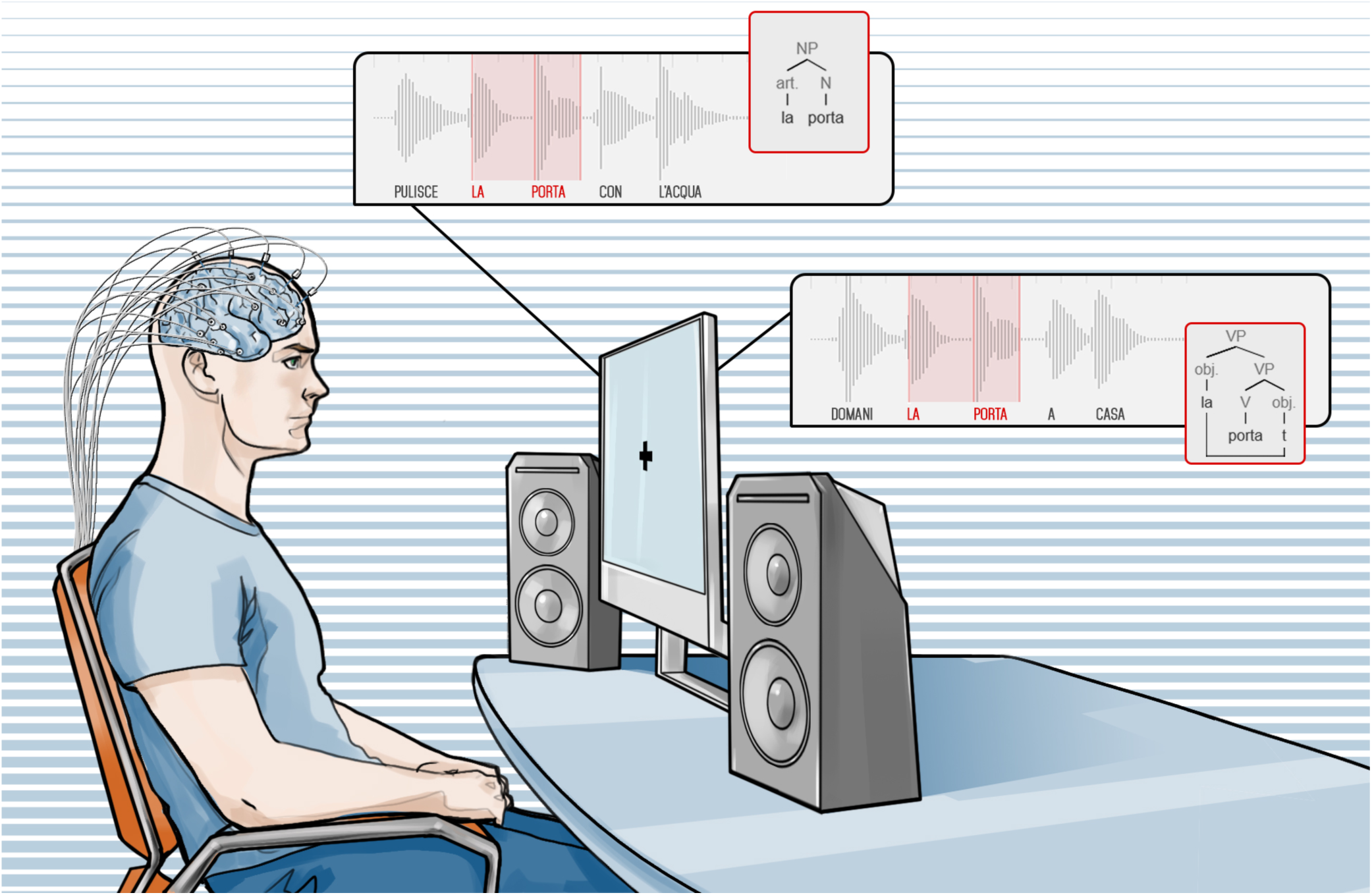
Example of auditory l stimuli presented to the subject. Languages may contain homophonous sequences. i.e. strings of words with the same sound and different syntactic structure. For example, in Italian, the very same sequence of phonems [la’pɔrta] may have two completely different meanings and two different syntactic structures: (i) LA (the) PORTA (door) as in PULISCE LA PORTA CON L’ACQUA (s/he cleans the door with water) (ii) LA (her) PORTA (brings) as in DOMANI LA PORTA A CASA (tomorrow s/he brings her home) In the first sequence [la’pɔrta] (written here as: *la porta*) is a Noun Phrase: the article *la* (the) precedes the noun *porta* (door). In the second sequence, instead, the very same sequence is a Verb Phrase: the object clitic pronoun *la* (her) precedes the verb *porta* (brings) which governs it. The difference is not only reflected in the distinct lexical classes, there is also a major syntactic difference: in the case of the noun phrase the element preceding the noun, namely the article, is base generated in that position; in the case of the verb phrase, instead, the element preceding the verb in the acoustic stimulus, namely the clitic pronoun, is based generated on the right of the verb occupying the canonical position of complements and then displaced to a preverbal position. This fundamental syntactic difference is represented in the syntactic trees in the picture: “t” indicates the position where the pronoun is base generated in the VP. Notably, to exclude phonological or prosodical factors which may distinguish the two types of phrases, in our experiment the exact copy of the pronunciation of one phrase replaced the other in either sentence in the acoustic stimuli. In other words, subjects heard the very same acoustic stimulus for each homophonous phrase.

**Figure 2.**
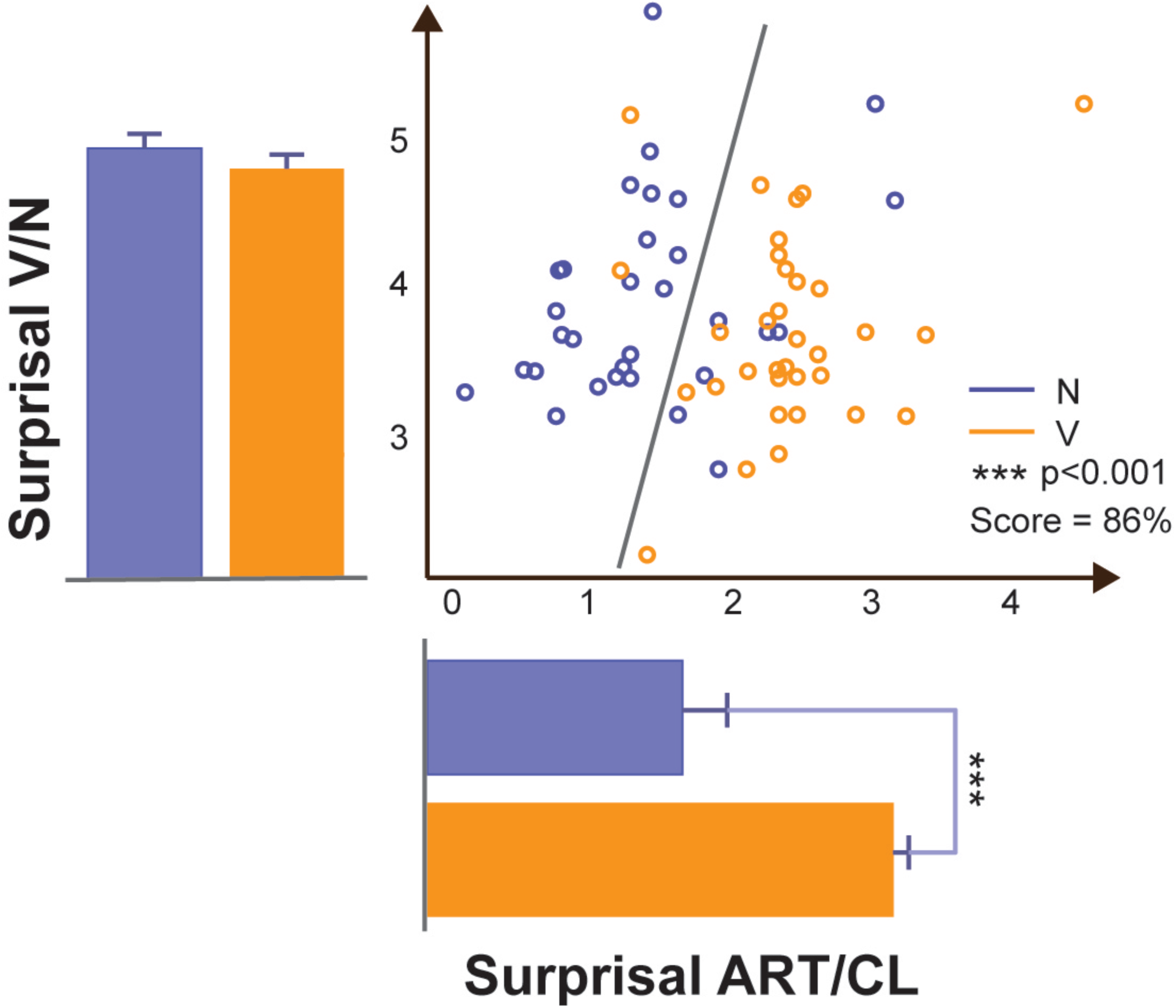
Surprisal analysis. Scatter plot of ITWAC surprisal values related to the Art/Cl (x axis) and Verb/Noun (y axis) position in the phrase. The gray line optimally separates (Support Vector Machine analysis) the Verb and Noun surprisal values with a Score of 86%. The bars represent the average and standard deviations of the surprisal values distribution for VPs (orange) and NPs (blue), tested for significance (ANOVA, 1 way). *** = p<0.001.

A total of 23 patients undergoing surgical implantation of electrodes for the treatment of refractory epilepsy^10^ completed all experimental sessions. Only patients without anatomical alterations, as evident on MR, were included. No seizure occurred, no alterations in the sleep/wake cycle were observed, and no additional pharmacological treatments were applied during the 24 h before the experimental recording. Neurological examination was unremarkable in all cases; in particular, no neuropsychological and language deficits were found in any patient. In all patients, language dominance was assessed with high frequency stimulation (50 Hz, 3 mA, 5 sec) during SEEG monitoring. Two patients also underwent a fMRI study during a language task before the electrodes implantation. Eight patients were excluded after analysis as they exhibited pathological EEG findings. Five patients were also excluded because no explored recording contact showed a task-related significant activation. Demographic data are shown in Extended Data Table 3. In the remaining 10 subjects, a total of 164 electrodes were implanted (median 16.5 range 13-19), corresponding to 2186 recording contacts (median 210; range 168-272). The number of contacts in the grey matter was 1439 (65.8%); 586 recording contacts in the language dominant hemisphere (DH). The DH was explored in 5 subjects (median electrodes 16, range 3-18; median contacts 210, range 25-225). The non-dominant hemisphere (NDH) was explored in 6 subjects (median electrodes 15, range 14-19; median contacts 208, range 182-272). SEEG exploration involved both hemispheres with a preference for the non-dominant side in 1 patient.

The temporal lobe was the most explored brain region, with 26 electrodes in DH and 42 electrodes in NDH, followed by frontal lobe (22 electrodes in DH and 21 in NDH).

The central lobe was implanted with a total of 22 electrodes (9 in DH). The Parieto-Occipital region was studied with a total of 9 electrodes in DH and 21 in NDH.

The contacts that exhibited a significantly different response according to whether the homophonous words belonged to VPs or NPs are considered “responsive contacts” (RCs). An example of RC is shown in Figure 3.

**Figure 3:**
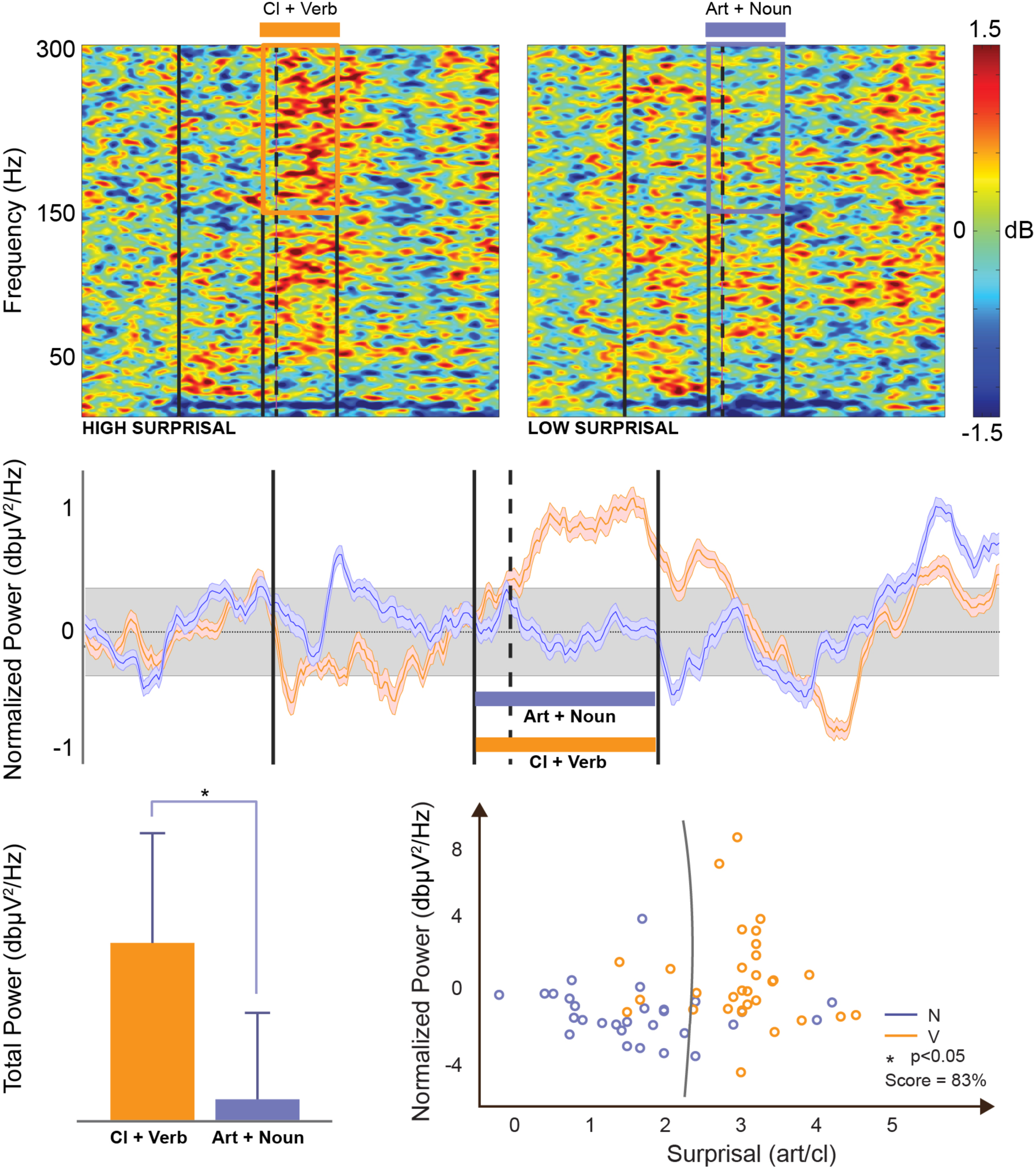
Example of event-related spectral perturbation and decoding for a responsive contact (channel). The first row represents the Event-Related Spectral Perturbation (ERSP) for VPs (left) and NPs (right) respectively. The four vertical lines respectively represent the beginning of the phrase, the beginning of the Art/Cl (homophonous phrase), the beginning of the first word after Art/Cl (i.e., Verb/Noun), the beginning of the word after that. The high gamma time Region of Interest (tROI) is highlighted by a superimposed square over the ERSP plots. The second row shows the baseline-normalized power in the [150 – 300] Hz spectrum interval. The four vertical bars have the same meaning as in the ERSP plots. The third row shows on the left a comparison between the VP tROI (orange) and NP tROI (blue) high gamma power. Similarly, the scatter plot on the right represents the tROI normalized power (y axis) and surprisal (x axis). The gray line optimally separates (Support Vector Machine analysis) the Verb and Noun classes in the surprisal/power feature space with a Score of 83%.

To validate the setup processing pipeline we analyzed the ERPImage and event related spectral perturbation (ERSP) of contacts responsive to the auditory stimuli (i.e., Heschl) and highlighted clear auditory event-related potentials (ERPs) and power increase time locked to the stimuli presentations (Extended Data Figure 2). Also, we retained in the RCs pools only the contacts where the different response between VPs and NPs was specific to the time region of interest (tROI, time interval that spans from the beginning of Art/Cl to the end of Noun/Verb). Incidentally, the high gamma frequency interval (150 – 300 Hz) showed the greatest tROI specificity in RCs. As an example, a RC (B13) is compared to a Heschl contact in Extended Data Figure 3. Only B13 shows (i) significantly higher power in the VP high gamma [150 – 300] time ROI (S) with respect to NP (Panel C, 4^th^ row, bottom right) and(ii) a significant power difference between VP and NP high gamma in the time ROI (ΔS) with respect to other time periods in the phrase, e.g., from the beginning of the phrase to Art/Cl, ΔA1 (Panel C, 4^th^ row, bottom left).

The ERSP analysis indicated that 242 (16.2%) of the leads exploring grey matter exhibited a significant high gamma (150Hz – 300Hz) power increase during the presentation of the corresponding VP homophonous phrase with respect to both the baseline and the other words (113 DH, 129 NDH).

The percentage of RCs in the DH was higher compared to that in the NDH (19.3% vs. 15.1%) and the difference was significant (p= 0.044, Fisher’s exact test).

The majority of RCs was found in the temporal lobe (133; 54.9%; 62DH; 71NDH), the majority of which in the middle temporal gyrus (55; 22.7%; 29 DH; 26 NDH) and in the superior temporal gyrus (9; 3.7%; 7 DH; 2 NDH). Out of 44 RCs (18.2%; 10 DH; 34 NDH) found in the frontal lobe, the majority of which were in the inferior frontal gyrus (13; 29.5%; 3 DH; 10 NDH) and in the frontal part of cingulate gyrus (20; 45,5%; 2 DH; 18 NDH). A detailed description of the localization of RCs for each patient can be found in Extended Data Table 4. Figure 4 shows all RCs positioned and template-matched after warping each patient’s MRI scan^11^.

**Figure 4:**
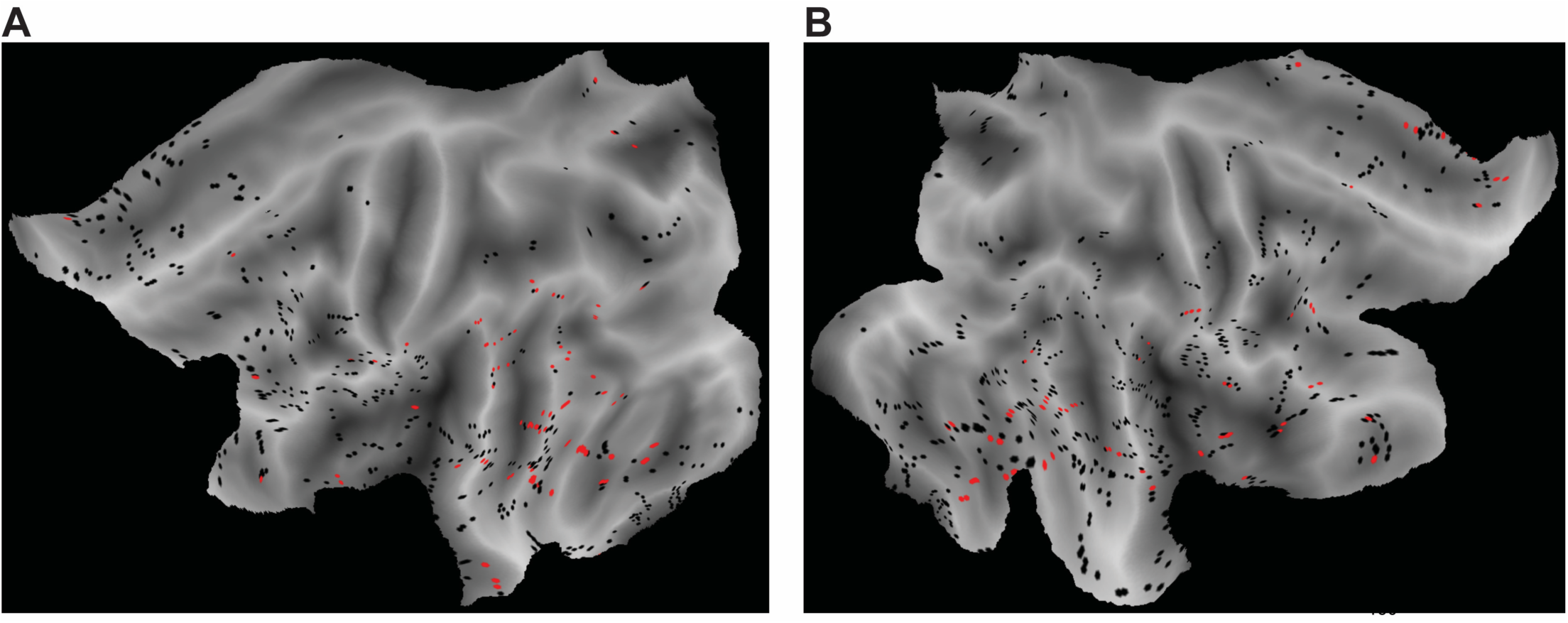
Main Responsive Contacts. Responsive contacts in the dominant (left panel) and non-dominant (right panel) hemispheres, merged across subjects over an average MRI template, group level.

Within the population as a whole, the significant difference was always due to higher ERSP for VPs with respect to NPs. High gamma activity (>100Hz) is receiving a growing interest to understand and characterize inter-regional cortical communications^12^. This band is one of the most used indices of cortical activity associated to cognitive function, and has been shown to be correlated with the neuronal spiking rate and to the hemodynamic BOLD response measured with functional magnetic resonance in both animal models^13^ and in human cortex^14, 15^. In particular, a large body of studies have indicated its value in tracking cortical activity during language perception and production^16^, supporting its use as a safer alternative to cortical stimulation for the presurgical mapping of cortical language areas^17^. In the present study, a significant increase of high gamma event related spectral perturbation^18^ (ERSP) was a specific index of the exposure to the syntactic contrast between clitic-verb phrases as compared to homophonous article-noun phrases.

This specific impact of syntactic structure on high gamma activity was not limited to the areas traditionally associated with syntactic processing on the basis of lesion effects and functional magnetic resonance evidence^19^. It must be underlined that a robust literature indicates that a categorial morphological contrast between Nouns and Verbs is indeed reflected in terms of lateralization and localisation^20^. The different activity observed in our experiment must then be related to a different factor, namely the syntactic structure of the stimuli. In particular, given that the physical stimuli where the same and that we did not observe the typical correlates distinguishing distinct lexical categories, such as noun and verbs, the higher activity of VPs can reasonably be correlated with the surviving difference, namely syntactic structure involving the operation of displacement of the object clitic from the right to the left side of the verb. Furthermore, these results suggest that, while syntactic impairment is known to be caused by focal lesions affecting nodal structures in a dedicated network, syntactic processing must involve a much more integrated pattern of brain activity than expected. Finally, our results concerning syntactic structures converge with parsing as shown by the surprisal analysis. Syntactic surprisal is related to the expectedness of a given word’s syntactic category given its preceding context^21^ and is associated with widespread bilateral activity indexed by the BOLD signal^22^. In fact, the position of the object to the left of the verb is reflected in the higher surprisal, showing that this measure is sensitive to syntactic structure.

All in all, the results found in confronting homophonous VPs and NPs allow us to factor out sound from the electrophysiological stimulus and consequently highlight a specific syntactic information distinguishing these universal linguistic structures. Notice that this separation could by no means be obtained by analyzing the electrophysiological correlates of inner speech since it has been proved that acoustic information is represented in higher language areas even when words are simply thought^5^. This first step provided here opens up to a deeper understanding of the structure and nature of human language and contributes to the ultimate far reaching goal of a complete neural decoding of linguistic structures from the brain^23^.

## Acknowledgements

We thank Elia Zanin for his help during the preparation of the stimuli. This work was partly funded by the Bertarelli Foundation, by the Ministero Istruzione Università e Ricerca (MIUR) DM 610/2017 and by the European Union’s Horizon 2020 research and innovation programme under Marie Skłodowska Curie grant agreement No. 750947 (project BIREHAB).

## Author Contributions

AM conceived the theoretical linguistic paradigm. FA, SC, SM, AM designed the experiment. AM construed the auditory stimuli. FA developed and integrated the experiment setup for stimuli presentation and synchronized recordings and developed the data processing algorithms. FA, PdO, VP, IS and GLR performed the recordings of the electrophysiological data. FA, IS, PdO analysed the data, critically reviewed the results at each analysis step and wrote the results. EC and FC elaborated the lexical and frequency statistics. FB recorded and edited the stimuli. SM supervised the processing activities. FA, SC, SM, AM wrote the paper. All the other authors provided comments to the manuscript and approved it.

## Competing interests

The authors do not have anything to disclose.

---

**Extended Data Figure 1:**
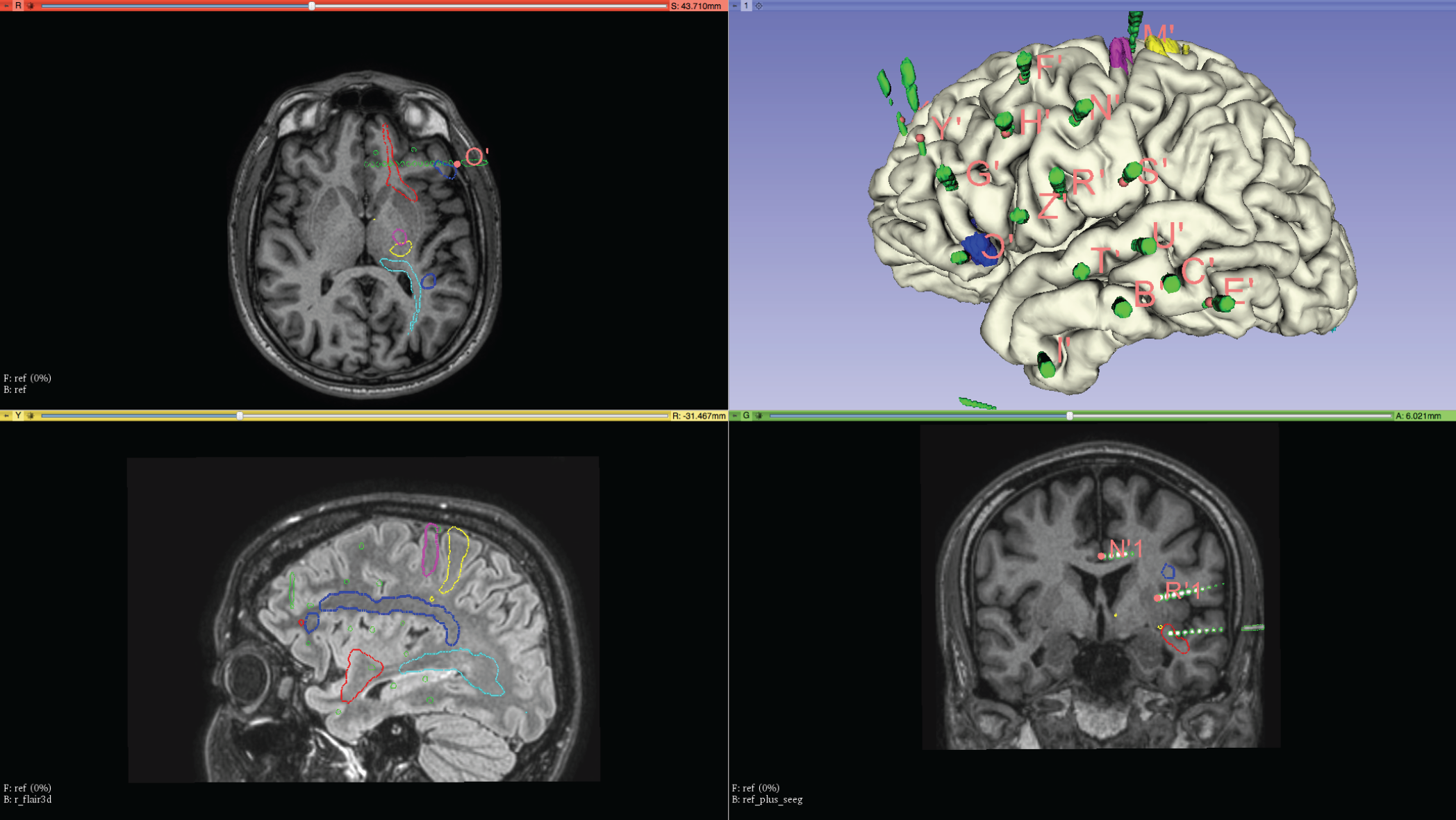
Example of Multimodal and interactive scene assembled with 3D Slicer software of patient s11.

**Extended Data Figure 2:**
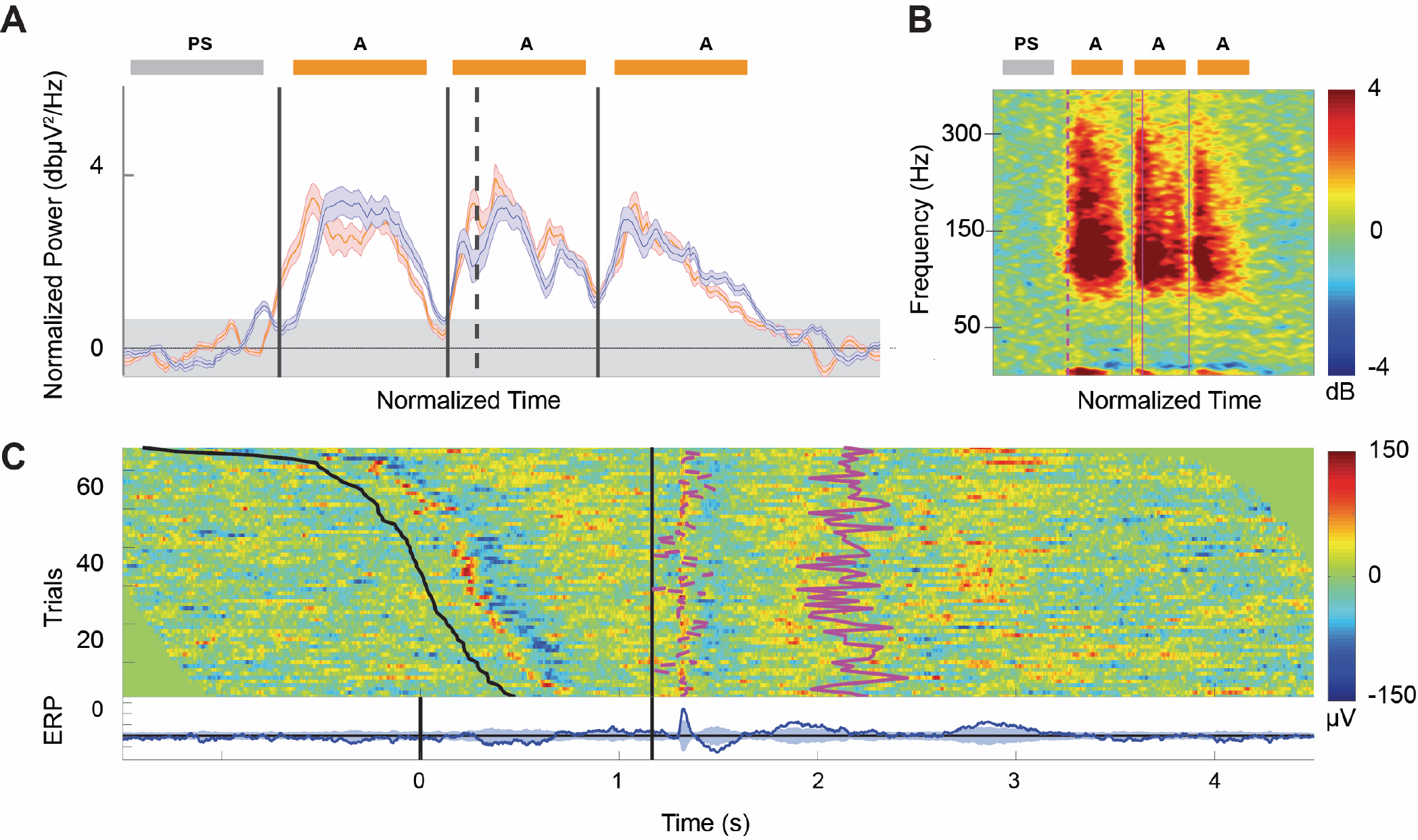
Heschl ERPImage and Time Frequency. **A)** Band-specific (150 – 300 Hz) ERSP (bERSP) for VPs (red) and NPs (blue) respectively for a Heschl contact. The four vertical bars separate the auditory stimuli (A) from the prestimulus interval (PS) and respectively indicate the beginning of the phrase, the beginning of the Art/Cl, the beginning of the word immediately following Art/Cl (Verb/Noun), the beginning of the word after. **B)** Time-warped Event-Related Spectral Perturbation (ERSP) of verb and noun phrases pooled together. The four vertical bars have the same meaning as in the baseline-normalized power plots (panel A). **C)** Event-related single-trial potential image (ERPImage) time-locked to the stimulus presentation. Trials are aligned to the beginning of Art/Cl (continuous black vertical line). The first black line indicates the beginning of the phrase for each trial. The following two pink lines respectively indicate the beginning of the word immediately following Art/Cl (Verb/Noun) and the beginning of the word after, for each trial.

**Extended Data Figure 3.**
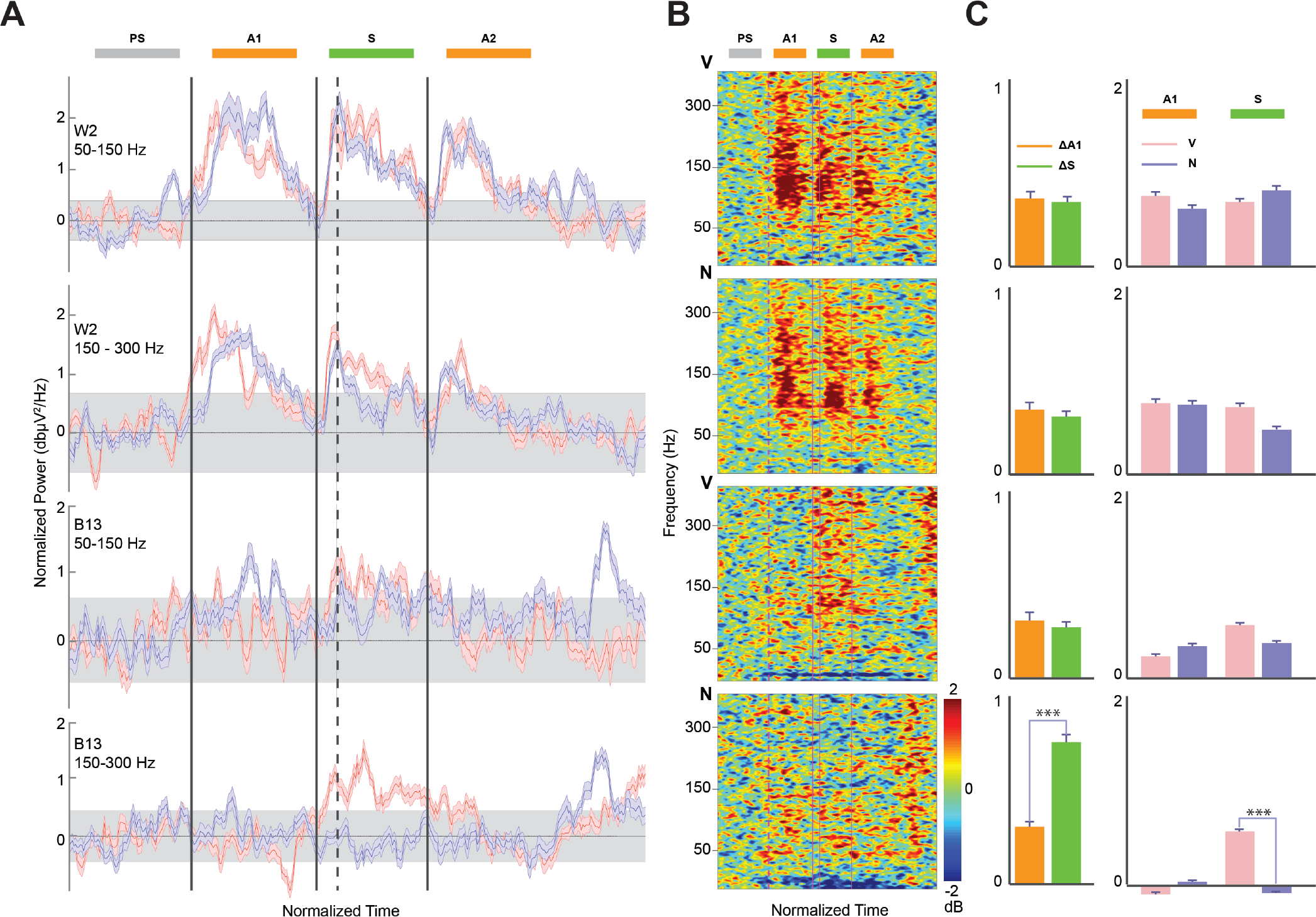
Band-specific ERSP relative to the [50 – 150] Hz and [150 – 300] Hz frequency intervals for a responsive contact (B13) and a non-responsive contact (Heschl – nonspecific auditory response). VPs are represented in pink, NPs in blue. The four vertical bars separate the regions pertaining to the auditory stimuli (A1, S, A2) from those within the pre-stimulus interval (PS). They respectively indicate the beginning of the phrase, the beginning of the Art/Cl, the beginning of the first word immediately following Art/Cl (Verb/Noun), the beginning of the word after. A1 and A2 refer to the auditory stimuli respectively before and after the tROI (Art/Cl + N/V syntax construct). **B)** Time-warped Event-Related Spectral Perturbation (ERSP) respectively for VPs and NPs. The four vertical bars and indicators have the same meaning as in the baseline-normalized power plots (panel A). **C)** From left to right the bars represent (i) the absolute value of the normalized power difference between VPs and NPs in the intervals A1 (orange) and S (green), (ii) the absolute value of the normalized power of VPs (pink) and NPs (blue) in the intervals A1 and S respectively.*** = p<0.001

**Extended Data Table 1:** surprisal values analyses. Number of valid cases, percentage of missing, mean and the standard deviation relative to the surprisal value, separately for the two experimental conditions (Noun-phrase, Verb-phrase).

|  | Noun-phrase |  |  |  | Verb-phrase |  |  |  |
| --- | --- | --- | --- | --- | --- | --- | --- | --- |
|  | Article |  |  |  | Pronoun |  |  |  |
|  | *valid* | *miss %* | *mean* | *std* | *valid* | *miss %* | *mean* | *std* |
| freq. Repubblica form | 30 | 3.23 | 1.45 | 0.50 | 21 | 32.26 | 2.95 | 0.52 |
| freq. ITWAC form | 31 | 0 | 1.46 | 0.48 | 22 | 29.03 | 2.84 | 0.64 |
| freq. Repubblica lemma | 30 | 3.23 | 1.14 | 0.86 | 31 | 0 | 2.23 | 0.66 |
| freq ITWAC lemma | 31 | 0 | 1.39 | 0.68 | 31 | 0 | 2.39 | 0.63 |
|  | Noun |  |  |  | Verb |  |  |  |
|  | *valid* | *miss %* | *mean* | *st.dev.* | *valid* | *Miss %* | *mean* | *st.dev.* |
| freq. Repubblica form | 31 | 0 | 3.90 | 0.76 | 18 | 41.94 | 3.73 | 0.9 |
| freq. ITWAC form | 31 | 0 | 3.78 | 0.57 | 23 | 25.81 | 4.07 | 1.16 |
| freq. Repubblica lemma | 31 | 0 | 3.67 | 1.09 | 31 | 0 | 3.80 | 0.71 |
| freq. ITWAC lemma | 31 | 0 | 3.90 | 0.72 | 31 | 0 | 3.72 | 0.71 |

**Extended Data Table 2:**
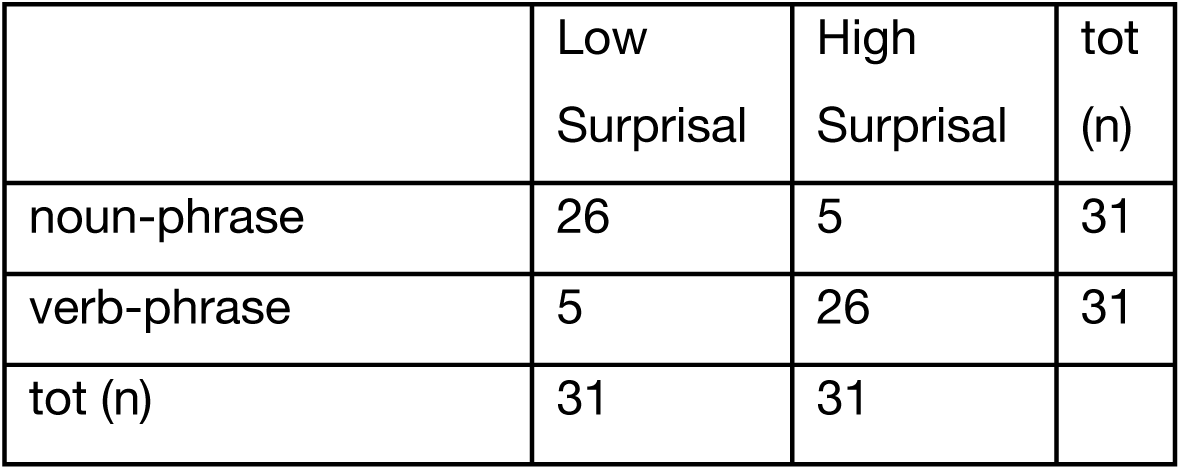
distribution of the surprisal values of articles and clitics on the basis of the median (M= 1.9097) obtained from the occurrence of the lemma in the ITWAC database.

|  | Low<br>Surprisal | High<br>Surprisal | tot<br>(n) |
| --- | --- | --- | --- |
| noun-phrase | 26 | 5 | 31 |
| verb-phrase | 5 | 26 | 31 |
| tot (n) | 31 | 31 |  |

**Extended Data Table 3:** Demographic data. Summary of demographic data of patients that successfully completed the study. SEEG: Stereo-Electro-Encephalography.

|  |  |
| --- | --- |
| **Number of patients** | **10** |
| **Gender** | **5 male and 5 female** |
| **Median age at SEEG** | **32 (range 17-44)** |
| **Median years of**<br>**scholarization** | **13 (range 0-17)** |
| **Median age of onset of**<br>**epilepsy** | **16 (range 2-30)** |
| **Median duration of epilepsy** | **15 (range 3-30)** |

**Extended Data Table 4:**
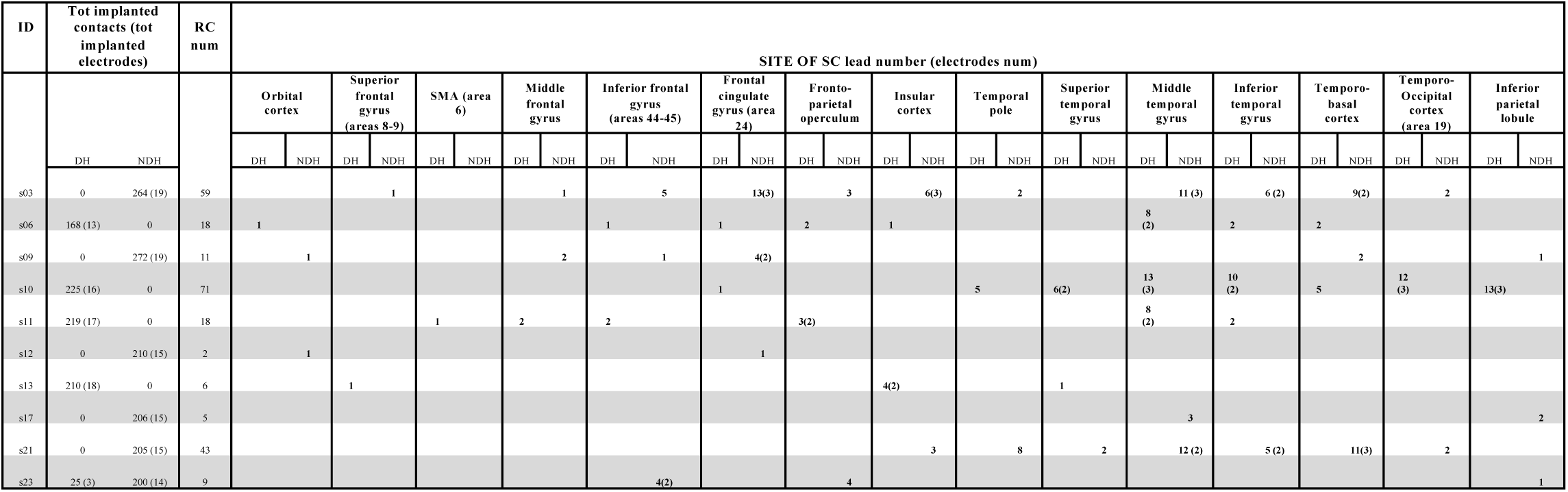
Responsive contacts. Highlights of responsive contacts in the whole population of patients that completed the study. RC: Responsive Contacts; SMA: Supplementary Motor Area; DH: Dominant Hemisphere; NDH: Non-Dominant Hemisphere. 6 leads (3 electrodes) in patient s10 showed an auditory response into the mesial cortex of occipital lobe.

## METHODS

### Stimuli

A novel set of stimuli which capitalizes on three special characteristics of Italian has been provided. First, some definite articles (such as [la] written as *la*; “the _fem.sing_.”) are pronounced exactly like some object clitic pronouns (such as [la] written as *la*; “her _fem.sing._”): both items are monosyllabic morphemes inflected by gender and number. Second, the syntax of articles and clitic pronouns is very different: like in English, articles precede nouns whereas complements follow verbs but, crucially, object clitics are obligatorily displaced the left of the verb with finite tenses. Third, the Italian lexicon contains several homophonous pairs of verb and nouns, such as [’pɔrta] (written *pɔrta*), which can either mean “door” or “brings”. Combining these facts together, a set of pairs of words such as [la ’pɔrta] (written as *la pɔrta*) has been construed which could be interpreted either as noun phrases (“the door”) or verb phrases (“brings it”) depending on the syntactic context (homophonous phrases) they are inserted in. Moreover, in order to be sure that no phonological or prosodical factors distinguish the two types of phrases, the exact copy of the pronunciation of one phrase replaced the other in either sentence in the acoustic stimuli. No other semantic or lexical distinction differentiated the two types of phrases which were balanced for major semantic features (such as abstract vs. concrete).

The acoustic stimuli were recorded using a Sennheiser Microphone MH40P48, Sound Card: Motu Ultralight Mk3, Connection: Firewire 400, Computer: Apple OSX 10.5.8. The stimuli were edited using Audiodesk 3.02 and mastered using Peak Pro7. Files were generated in 16bit, 44.1 kHz (Sampling Frequency); intensity was normalized to 0 Db and rendered in .wav format. All sentences were read by the same person: Italian native speaker, male, 53 years old.

### Surprisal value computation

The value of surprisal (S) indicates how unexpected a given word is on the bases of the preceding words^1^. In order to calculate the surprisal value associated to each word of the sentence, we used the algorithms developed by Roark^2^ with a model of Probabilistic Context Free Grammar 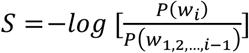, where *P*(*w*_*i*_) corresponds to the probability of occurrence of the target word and *P*(*w*_1,2,…,*i*−1_) to the probability of occurrence of the preceding words. More in details, given the word *w*_*i*_, the value of surprisal has been obtained through the logarithm of the ratio between the probability of occurrence of the bigram containing the target word (i.e., *Bigram_w*_*i*_) and the probability of occurrence of the word immediately preceding the target word (i.e., *Unigram_w*_*i*−1_). The formula is as follows: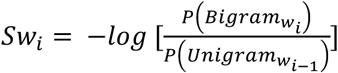. For instance, in the sentence *pulisce la porta con l’acqua* (“s/he cleans the door with water”) the value of surprisal associated with the Italian word *la* (the definite article preceding the noun) is: 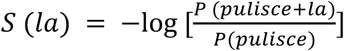

In order to obtain the frequency of unigrams (i.e., single words) and bigrams (i.e., pairs of words), we initially considered the online databases “La Repubblica”, a corpus derived from Italian newspaper texts written between 1985 and 2000 and containing about 380 million words and of the Italian WEB Corpus (ITWAC^3^, obtained from Italian texts on the Internet, composed by about 1.5 billion words. For each unigram and bigram, we reported both the occurrence of the form, i.e. word’s frequency considering the specific category (e.g. article vs pronoun), and of the lemma, i.e. word’s frequency without taking into account the differentiation into categories. We then calculated the value of surprisal (both derived from the occurrence of the form and the lemma, for both databases) associated with all the elements of the sentences belonging to the two experimental conditions (i.e., clitic + verb and article + noun).

Of note, sentences n.4 and n.63 were excluded from the analyses since they were not organized with the same structure as the other stimuli, i.e. article + noun or clitic + verb. Sentences n.51 and n.52 were excluded because both belonged to clitic + verb condition, namely containing the same target word. The data discussed were therefore related to 31 verb-phrases and 31 noun-phrases.

Statistical analyses were performed both on the form and on the lemma for both databases. Due to the presence of missing values (see the Table for the percentages of misses, divided by condition) we considered only the analyses based on ITWAC lemma values.

We compared the value of surprisal associated to the two target elements of the two types of experimental phrases by means of paired samples t-test. Specifically, we compared the value of surprisal associated with the article with that associated with the pronoun and the value of surprisal associated with the noun with that associated with the verb. Since sentences n.4 and n.63 were excluded (see previous paragraphs), the corresponding sentences, respectively, noun-phrase n.3 and verb-phrase n.64 were considered as the two paired elements in the analysis.

The analyses showed significant differences between the value of surprisal of the article compared to that of the pronoun (t= −6.794, p< .001), with a higher surprisal value found for pronouns than for articles. No significant differences were found between the value of surprisal associated to nouns and that associated to verbs (t= 1.357, p = .185).

In order to dichotomize the surprisal variable, we divided the distribution of the surprisal values of articles and clitics on the basis of the median (M = 1.9097) obtained from the occurrence of the lemma in the ITWAC database. The values were divided, respectively, into high and low surprisal.

### Patients

A total of 23 patients were recruited for the present study among those who underwent on surgical implantation of multi leads intracerebral electrodes for refractory epilepsy in the “Claudio Munari” Epilepsy Surgery Center of Milan in Italy^4,5^. Only patients with negative MRI and with no neurological and/or neuropsychological deficits were included. Based on anatomo–electro–clinical correlations, each patient-specific strategy of implantation was defined purely on clinical needs, in order to define the 3D shape of the epileptogenic zone (EZ).

Demographic data of the 10 patients included in the analysis are reported in Extended Data Table 3. The present study received the approval of the Ethics Committee of ASST Grande Ospedale Metropolitano Niguarda (ID 939-2.12.2013) and informed consent was obtained.

### Surgical Procedure and Recording Equipment

All trajectories of patient-related implantation strategy are planned on 3D multimodal imaging and the electrodes are stereotactically implanted with robotic assistance. The whole workflow was detailed elsewhere^6^. SEEG electrodes are probes with a diameter of 0.8 mm, comprising 5 to 18 2 mm long leads, 1.5 mm apart. A post-implantation Cone-Beam-CT, obtained with the O-arm scanner (Medtronic, Minneapolis, Minnesota), is subsequently registered to pre-implantation 3D T1W MR, in order to assess accurately the position of every recording lead. Finally, a multimodal scene is assembled with 3D Slicer^7^, aimed at providing the epileptologist with interactive images for the best assessment of anatomical electrical sources (Extended Data Figure 1).

During the experiment the SEEG was continuously sampled at 1000Hz (patients 1-12) and 2000Hz (patients 12-23) by means of a 192 channels SEEG device (EEG-1200 Neurofax, Nihon Kohden). In each patient, all leads from all electrodes were referenced to two contiguous leads in the white matter, in which electrical stimulations did not produce any subjective or objective manifestation (neutral reference).

### Recording Protocol

Each subject rested in a comfortable armchair. Constant feedback was sought from the patient to ensure the overall comfort of the setup for the whole duration experiment. Stimuli were delivered in the auditory modality (see also Figure 1) using Presentation from Neurobehavioral Systems software. Phrases were delivered via audio amplifiers at a comfortable volume for the subject (minimum volume for words to be perceived with ease, according to the subject) while gazing at a little cross on a screen (27 inches). A synchronization TTL trigger spike was sent to the SEEG trigger port at the beginning of auditory presentation (sentence). Jitter and delays were tested and verified to be negligible (less than 1ms). The whole experiment lasted around 30 minutes to maximize engagement. At the end of each task, subjects were asked to answer a few short questions on the content of the stimuli. Indeed, patients were always able to provide correct answers to the questions, thus demonstrating their continuous engagement to the task.

A camera, synchronized to the SEEG recording at source, was used to control for excessive blinking, maintenance of fixation with no eye movement, silence and any unexpected behavior from the patients.

### Control Experiment

As a further control for the analysis, the first three subjects underwent an extra auditory task. The modalities remained the same, however the sounds were substituted with beeps (auditory presentation) not carrying any meaning at all.

### Data analysis

A band-pass filter (0.015–500 Hz) applied at hardware level prevented any aliasing effect from altering SEEG data. Recordings were visually inspected by clinicians and scientists in order to ensure the absence of artifacts or any pathological interictal activity. Pathological channels were discarded. Further analyses were carried out using custom routines based on Matlab, Python and the EEGlab toolbox^29^. Data were annotated with the events triggered by the beginning of each stimulus. Events were time locked to the beginning of each word (initial syllable of the word for auditory presentation). Epochs were extracted in the intervals [-1.5 4.5] s, time-locked to the initial presentation (i.e., beginning of the phrase). The length of the epoch was selected so as to always include the complete stimulus presentation (trial). Epochs with prominent artifacts (e.g. spikes) over significant channels were rejected. To determine significant responsive sites, analyses were performed both in the time and frequency domains. Epochs were then sorted into two classes based on the surprisal value (low or high).

### Analyses in the time domain

In the time domain, single-trial data epochs were color-coded by amplitude to form a ERPImage 2D view^8^, without any smoothing over trials (Extendend Data Figure 2, panel C). The ERPImage allowed to assess the presence of Event-Related Potentials (ERPs) and their significance over time (e.g., to verify the presence of any habituation phenomena) with respect to the baseline. The ERPImage analysis was performed both (i) after time-warping the trials so as to temporally align the other events and (ii) after aligning the trials to the beginning of the beginning of Art/Cl position in the phrase and annotating the relative position of the other events (i.e., beginning of the sentence, beginning of the first and second words after Art/Cl).

### Analyses in the frequency domain

Time-frequency transforms of each trial were normalized to the baseline (divisive baseline, ranging from-1500ms to -5ms time-locked to the beginning of the sentence), time-warped to the beginning of the sentence, beginning of Art/Cl, beginning of the first and second word after, then they were averaged across trials to obtain the event-related spectral perturbations (ERSPs) a generalization of ERD/ERS analyses to a wider range of frequencies^30^ (Ext. Data Figure 2, Panel B), i.e., from theta (1-4Hz) to high gamma (150-300 Hz). A bootstrap distribution over the trials baseline was used to determine significance (p<0.05) of the time-frequency voxels. We considered the average ERSP across the Gamma ([50 – 150] Hz) and High Gamma ([150 – 300 Hz]) frequency bands to obtain band-specific ERSP (bERSP) and compared it over time between low and high suprisal (Ext. Data Figure 2, Panel A). These bands were selected after a preliminary analysis of data related to Heschl gyrus in real and control experiments, which highlighted the presence of significant bERSP up to 300 Hz (see Ext. Data Figure 2 panel B). The preliminary analysis also showed that several contacts reported a significant time-specific differentiation in high gamma ([150 – 300] Hz) bERSP between VPs and NPs and we used that frequency band to highlight responsive contacts (see next paragraph).

### Identification of responsive contacts

Each contact (i.e., channel) for each subject underwent a series of screenings to determine its significance. A contact was deemed responsive if either low or high surprisal high gamma bERSP had significant amplitude specifically in the tROI (interval that spans from the beginning of Art/Cl to the end of Noun/Verb), for a significant time span. The amplitude was deemed significant if and only if greater than 95% of the distribution of amplitudes across frequencies for a significant time span. A time span was deemed significant if longer than the 95% of significant intervals in the baseline. The rationale of this test was to exclude those contacts that did not reach significance in the time ROI and ensure specificity in frequency (i.e., statistically different low and high surprisal high gamma time courses – only one of them being over threshold, or both being over threshold but statistically different – p<0.05), and time, (i.e., no significance when performing the same analysis at other time intervals such as from the second word after Art/Cl to the end of the sentence or from the beginning of the sentence to the beginning of the Art/Cl). Significant contacts were then ranked from high to low *sig* values according to the formula *sig = a* * *t/* ∑ *a*_*i*_ *t*_*i*_ where *a* is the maximum amplitude over the time ROI, *t* is the length of the interval within the time ROI the amplitude is significant, *a*_*i*_ and *t*_*i*_ respectively the maximum amplitude and length of the interval at the other positions (i) in the phrase (i.e., outside the tROI). The rationale of this formula was to determine the contacts that highlighted the maximum time-specific significant difference.

An inspection of all the contacts was also visually performed by expert clinicians and results were compared to the data-driven analysis in a double-blind fashion. The concordance was 84%. This analysis provided both validation to the data-driven analysis and also provided an extra control that selected responsive contacts (i) were not located in the white matter, (ii) were not located in affected regions of the brain, (iii) exhibited similar behaviour (e.g., high gamma time course waveform shape) if anatomically close and referring to the same brain region.

### Decoding

Decoding of the phrase type (noun and verb phrases) was first performed based on the surprisal relative to the Art/Cl and Verb/Noun parts of the phrases (Fig. 2). After testing for normality (Kolmogorov-Smirnov), VP and NP surprisal values were also statistically compared (ANOVA, 1-Way). Decoding of the two classes was also performed on the feature space formed, for each trial, by the Art/Cl surprisal value and the power amplitude in the time ROI (Fig.3). In both cases a Support Vector Machine (SVM) algorithm with leave-one-out cross validation (LOOCV) was implemented to ensure the generalizability of the model.

